# Review: Assessing available genetic diversity estimates of rare breeds of livestock

**DOI:** 10.1101/2025.05.14.653764

**Authors:** C. M. Rochus, C. F. Price, I. Pocrnić

## Abstract

The United Kingdom is home to many local and rare livestock breeds. The local breed populations are highly adapted to specific environments in the United Kingdom and these and other rare breeds provide solutions to niche needs. Rare breeds often contribute more than expected to overall species genetic diversity, which is important because genetic variation is needed for adapting to new challenges. It is therefore very important from both United Kingdom and global perspectives to maintain genetic diversity of rare livestock breeds in the United Kingdom, and to do this, we need to evaluate the monitoring of genetic diversity to identify gaps in our knowledge and prioritise resources reserved for conservation purposes. The objectives of this study were to survey the literature to: (1) summarise genetic/genomic characterisation (effective population size (**N_e_**) and inbreeding) of domestic populations (livestock and equine) in the United Kingdom and Ireland; (2) compare number of populations on the United Kingdom’s Rare Breeds Survival Trust watch list and the number of United Kingdom and Irish populations in the peer-reviewed literature with inbreeding, genetic diversity and/or N_e_ estimates; and (3) compare annually reported (census-based) estimates of N_e_ with (inbreeding- and DNA-based) estimates from peer-reviewed literature. We found a total of 37 publications with N_e_ or inbreeding estimates for United Kingdom or Ireland livestock populations, published from 1975 to 2024. While many (42%) of the breeds on the Rare Breeds Survival Trust watchlist have been included in publications, there are still many breeds, and a few species (turkey, duck and geese) with no publicly available pedigree- or DNA-based genetic diversity measures. We found census-based N_e_ estimates were, on average, higher than DNA-based estimates, likely due to violated assumptions when estimating N_e_ with census-based data because of the way livestock mating systems are designed. Most peer-reviewed papers estimated genetic diversity measures using pedigree, microsatellite markers, or single nucleotide polymorphisms (**SNP**) markers. To identify breed-unique variants responsible for adaptive traits in rare breeds, more studies using whole-genome sequencing will be needed. Altogether, we have summarised genetic diversity estimates of United Kingdom livestock populations, identifying gaps in knowledge.

**Implications:** Maintaining genetic diversity in livestock helps safeguard against future challenges to our food production systems. We surveyed publicly available measures of genetic diversity for farm animal populations in the United Kingdom (including government, Rare Breeds Survival Trust, and peer-reviewed scientific publications) to identify any gaps. We found census-based estimates of genetic diversity, which are used to inform and prioritise resources for conservation efforts, on average, are likely overestimating available genetic diversity. Additionally, most farm animal breeds had only these census-based estimates. Genetic diversity estimates could be improved using pedigree or DNA information, more accurately identifying endangered breeds.

## Introduction

Livestock populations in the United Kingdom originated in centres of domestication in Southwest Asia, East Asia, and Mesoamerica, with the first domesticated species arriving to the British Isles during the Neolithic transition to agriculture (Larson et al., 2014; Whittle et al., 2011). Through this process, livestock populations have become highly adapted to many niche environments (Zeder, 2008). Those populations today are called local breeds and are often maintained by breed associations for their unique qualities and nostalgia. These breeds can be described as hardy, many of which can thrive on low-quality feed not appropriate for human consumption and habitat in areas and weather conditions where other breeds cannot. Rare breeds have lost their historic popularity because commercial breeds are more productive (Canali, 2006). As a consequence of decreasing population sizes, rare breed societies do not have the resources to support breeding programmes, unlike contemporary commercial populations that can use genomic selection to inform mating decisions (DEFRA, 2021). However, there are several roles that rare breeds fill in the modern world. In the United Kingdom and European Union, niche products, like cheese and meat, from local breeds are sold with a geographical indicator (protected designation of origin, and protected geographical indication) (‘Geographical indications and quality schemes explained’, n.d.; ‘Protected geographical food and drink names: UK GI schemes’, 2024). Local breeds in the United Kingdom are used for conservation grazing to maintain numerous habitats (grassland, heathland, wood pasture, coastal and floodplains, fen, scrub and scrub mosaics, saltmarsh and sand dunes habitats) (‘Graze with livestock to maintain and improve habitats’, n.d.) and are used in organic farming (Van Diepen et al., 2007).

Local breeds also play an intangible cultural role in their communities providing a glimpse into the past, including domestication processes and the Neolithic transition to agriculture, all the way up to modern times. Finally, in an ever-changing world, these historic breeds can play an important role as reservoirs of genetic variation, providing the variation needed for sustainable agriculture systems with animals that can adapt to future challenges, which has been recognised by the United Nations (UN environment programme, 2022; United Nations General Assembly, 2015).

There are many other important roles for local breeds to play in modern society, besides maintaining total livestock species genetic diversity. To highlight a few examples from the United Kingdom, we present breeds with unique traits and their current and historic uses. The North Ronaldsay sheep, found primarily in the Orkneys has adapted to primarily eat seaweed, mainly composed of brown kelp, after they were mostly contained to the shore in the 19th century to protect cultivated land from being grazed (RBST, n.d.; Porter, 2002). Many modern breeds struggle with some functional traits as a result of a focus on improving production traits (Rauw et al., 1998). In contrast, local breeds are prized for these functional traits, and Shetland cattle are an example of this with their excellent calving ease (Schwabe and Hall, 1989), they can be bred with larger continental breeds without calving issues (RBST, n.d.). Local breeds can provide unique animal products that cannot be provided by commercial breeds. One of the first protected animal products in the European Union was the Gloucestershire Old Spots pig (RBST, n.d.). In the application for this status, it was indicated that texture, and flavour of these pigs are distinguishable from commercial pigs (The Gloucestershire Old Spots Pig Breeders’ Club, 2006). There were some traditional functions of local breeds that may not be in their current usage, but could inspire more sustainable practices. Before the use of pesticides to control parasites Shetland geese were traditionally grazed in pastures after sheep were grazed to manage parasites like liver fluke (RBST, n.d.). Finally, many of the local breeds can thrive and forage in any climate, like the Scots Grey, a traditional Scottish chicken breed traced back to the 16th century (RBST, n.d.). This ability to thrive outdoors is particularly useful for conservation grazing. Many iconic landscapes in the United Kingdom, like meadows and grasslands that harbour a third of British flora, were created from traditional farming practices (Peach, 2019). In the Great Trossachs Forest, grazing allows for a more diverse environment for wild birds, and opportunities for less competitive grass species to seed and germinate, among other benefits (The Great Trossachs Forest, 2025). They use a combination of Luing cattle, Hebridean sheep and Scottish Blackface sheep, each providing different grazing patterns to contribute to the park’s diversity (The Great Trossachs Forest, 2025).

Local breeds from the British Isles have played an outsized role in global commercial agriculture today. Breeds from the United Kingdom have had widespread popularity, for example, purebred and crossbred Angus, Jersey, and Guernsey cattle can be found in many countries around the world, many global commercial pig companies have developed lines based on Large White and Yorkshire breeds, and the Leicester Longwool was widely used in crossbreeding schemes to improve the carcass and fleece of sheep breeds in Europe, Australia, Canada, New Zealand, and the US (Porter, 2002). The Leicester Longwool was the product of a new type of breeding, “in-and-in”, developed by English farmer Robert Bakewell in the 18th century, where inbreeding was used to fix traits in a population (Wykes, 2004). This is believed to be the first implementation of systematic selective breeding, when in comparison, many famers to this point had been randomly mating livestock (Wykes, 2004).

The United Kingdom government recognises the importance of maintaining the genetic diversity of its local breeds and has several national and international obligations to conserve rare breeds, including the 2011 England Biodiversity Strategy, the United Nation’s Food and Agriculture Organisation’s “Global Plan of Action”, the United Nation’s Convention on Biological Diversity “Strategic Plan for Biodiversity 2011-2020”, and for input to the European Farm Animal Biodiversity Information System and Food and Agriculture Organisation’s Global Information System (Beale, 2024). To fulfil these obligations, the Department for Environment, Food and Rural Affairs in the United Kingdom monitors the number of active sires and dams for livestock populations by breed to inform and prioritise resources for conservation efforts (with data for cattle, equine, sheep, goat and pig publically available as far back as 2000, and for poultry available from 2021) (Beale, 2024).

The Department for Environment, Food and Rural Affairs collects this census data through collaboration with breed organisations and the Rare Breeds Survival Trust, a charity established in 1973 that conserves and promotes native livestock and equine breeds in the United Kingdom (RBST, n.d.). Census data is used to estimate effective population size (**N_e_**), which is the number of individuals in an idealised population that would generate the same rate of inbreeding as found in the population of interest (Falconer and Mackay, 1996), allowing for interpretation and comparison of N_e_ values across populations and species.

Additionally, several research groups have studied the genetic diversity and inbreeding of local and rare breeds, publishing their results in peer-reviewed literature. These studies were standalone projects or part of global consortia (Bovine HapMap (The Bovine HapMap Consortium et al., 2009), Sheep Hapmap, Adaptmap (for goats) (Stella et al., 2018), etc.). The objectives of this study were to survey the literature to: (1) summarise genetic/genomic characterisation (genetic diversity, N_e_ and inbreeding) of domestic populations (livestock and equine) in the United Kingdom and Ireland; (2) compare number of populations on the Rare Breeds Survival Trust list and the number of populations in the United Kingdom and Ireland in the peer-reviewed literature with inbreeding, genetic diversity and/or N_e_ estimates; and (3) compare annually reported (census-based) estimates of N_e_ with (DNA-based) estimates from peer-reviewed literature. By systematically reviewing the peer-reviewed genetic diversity literature for livestock populations in the United Kingdom, we can identify gaps, which will help with prioritising research areas.

## Material and methods

We used Web of Science (https://www.webofscience.com) to search (on 2025-01-13) using the following search term:

(inbreeding OR diversity OR “effective population size”) AND (cattle OR cow* OR bull* OR pig* OR chicken* OR sheep* OR goat* OR horse* OR duck* OR turkey* OR geese) AND (”UK” OR “United Kingdom” OR “Britain” OR “Scotland” OR “England” OR “Wales” OR “Northern Ireland” OR “Ireland”) AND (”genomic” or “genetic” or “pedigree”).

We searched for inbreeding, diversity and N_e_ estimates for all species identified on the Rare Breeds Survival Trust’s 2024-2025 watchlist for breeds in the United Kingdom (‘RBST watchlist 2024’, 2024). Irish populations were included due to the shared border between Northern Ireland and Ireland, making some livestock and equine populations to likely be well-connected. Values were extracted from articles, tables, and figures (using WebPlotDigitizer (Rohatgi, 2024)). The Rare Breeds Survival Trust watchlist (‘RBST watchlist 2024’, 2024) is annually published and used to prioritise breeds for conservation by considering genetic diversity and number of breeding females.

Every year the Department for Environment, Food and Rural Affairs collects census numbers (they can be number of animals registered or an estimate of total number) from breed organisations and the Rare Breeds Survival Trust, and publishes them (Beale, 2024). We have used the 2023 release of census and effective population estimates (Beale, 2024). This data is used to monitor numbers of animals over the years to help with prioritisation of conservation efforts. The Department for Environment, Food and Rural Affairs uses the following formula to estimate N_e_:

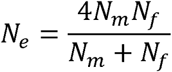

Where N_m_ is the number of breeding males, and N_f_ is the number of breeding females) (Wright, 1931). N_e_ is the number of individuals that would be needed for the inbreeding estimate to be true in an idealised population (Falconer and Mackay, 1996). An idealised population has the following conditions: (1) mating is restricted to a population with no migration, (2) generations are discrete, (3) number of breeding individuals are the same for every generation, (4) mating is random and includes self-fertilisation, (5) there is no selection, and (6) mutation is disregarded (Falconer and Mackay, 1996). The effective population estimate is standardised and comparable between populations, and can be used to estimate the rate of inbreeding (Wright, 1931). The United Nation’s Food and Agriculture Organisation’s guidance indicates N_e_ should be 50 animals per generation, or more, as an N_e_ of 50 corresponds to 1% inbreeding rate per generation (when number of breeding males and females are equal) (FAO Commission on Genetic Resources for Food and Agriculture, 2015).

### Study design

We have summarised the peer-reviewed literature on genetic diversity in livestock populations in the United Kingdom by counting the number of published papers per species, population, data type (pedigree, mitochondrial DNA, microsatellite, single nucleotide polymorphisms (**SNP**), or whole genome sequencing) and number of populations both in the literature and on the Rare Breeds Survival Trust 2024 watchlist.

Additionally, for all inbreeding- and DNA-based N_e_ estimates found for populations in the United Kingdom in the literature, we compared this with census-based N_e_ estimates published by the Department for Environment, Food and Rural Affairs. To make the closest comparisons, we compared census-based estimates from the same year of DNA sampling that was used in the DNA-based estimates. If the year of sampling was not available, year of publication was used as a proxy. If there was no census-based estimate for the year of sampling/year of publication of DNA-based estimates, the closest year with an estimate was used. If the sampling occurred over a period of years, census-based N_e_ estimates from those years were averaged. We used a paired t-test to determine if census-based and DNA-based N_e_ estimates were significantly different.

## Results

### Summary of literature search

In total, we have included 36 peer-reviewed publications and one report written for the Rare Breeds Survival Trust, published between 1975 and 2024, all of which have estimates of genetic diversity, including inbreeding coefficients, nucleotide diversity or N_e_ of livestock or equine populations in the United Kingdom or Ireland (see **Supplementary Table S1** for information on publications and diversity estimates included in this study). The 37 publications are summarised in **Table 1**, including sources of data for each study (original data from a total of 50 publications). **Figure 1** shows our workflow, including total number of papers found through the Web of Science search, number of papers removed due to not meeting certain criteria, and number of papers included that were not identified in the Web of Science search.

**Figure 1:**
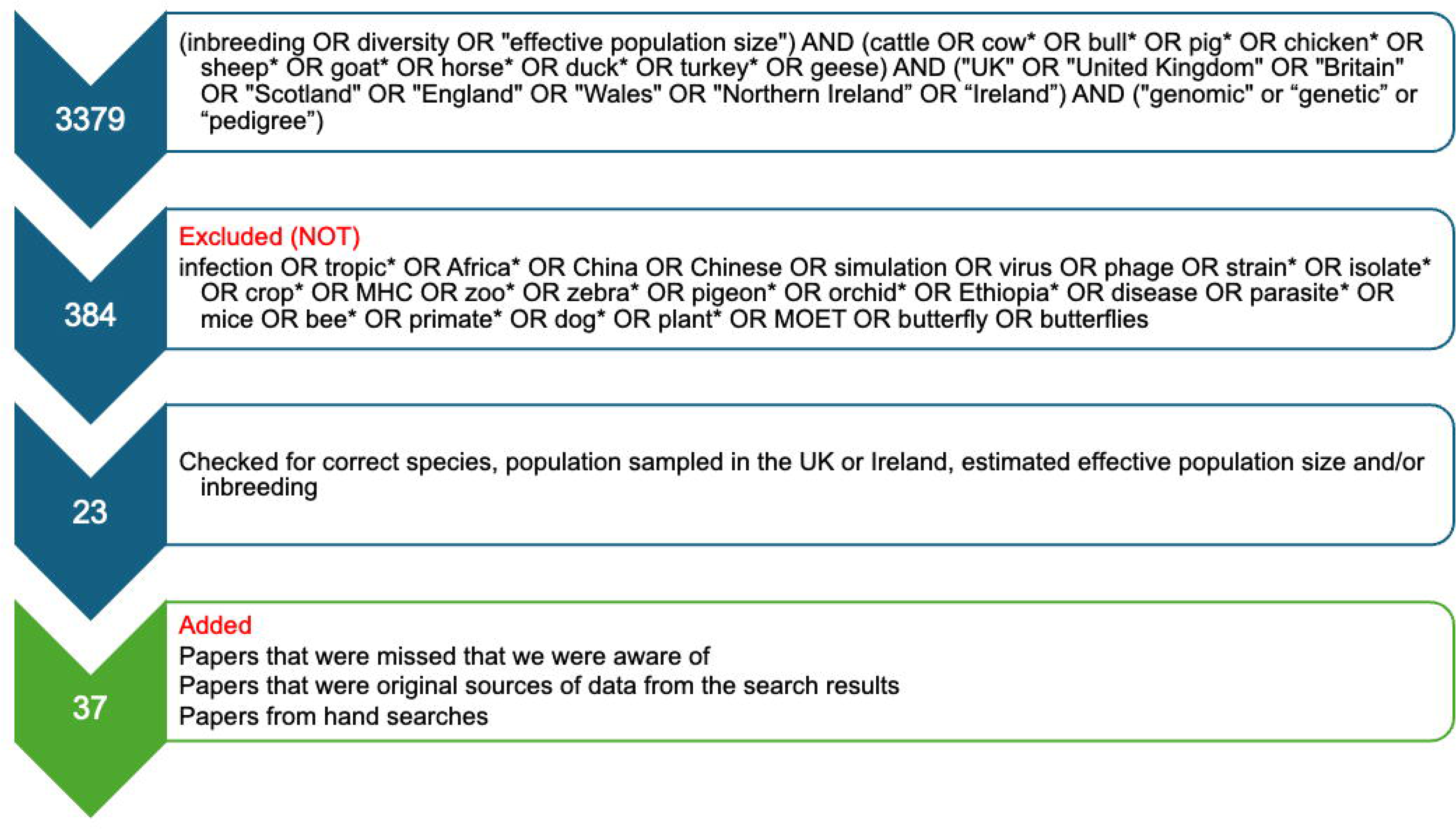
Literature search terms and number of results from Web of Science, for a search of genetic diversity measures of cattle, chicken, duck, equine, geese, goat, pig, sheep, and turkey populations in the United Kingdom and Ireland.

**Table 1:**
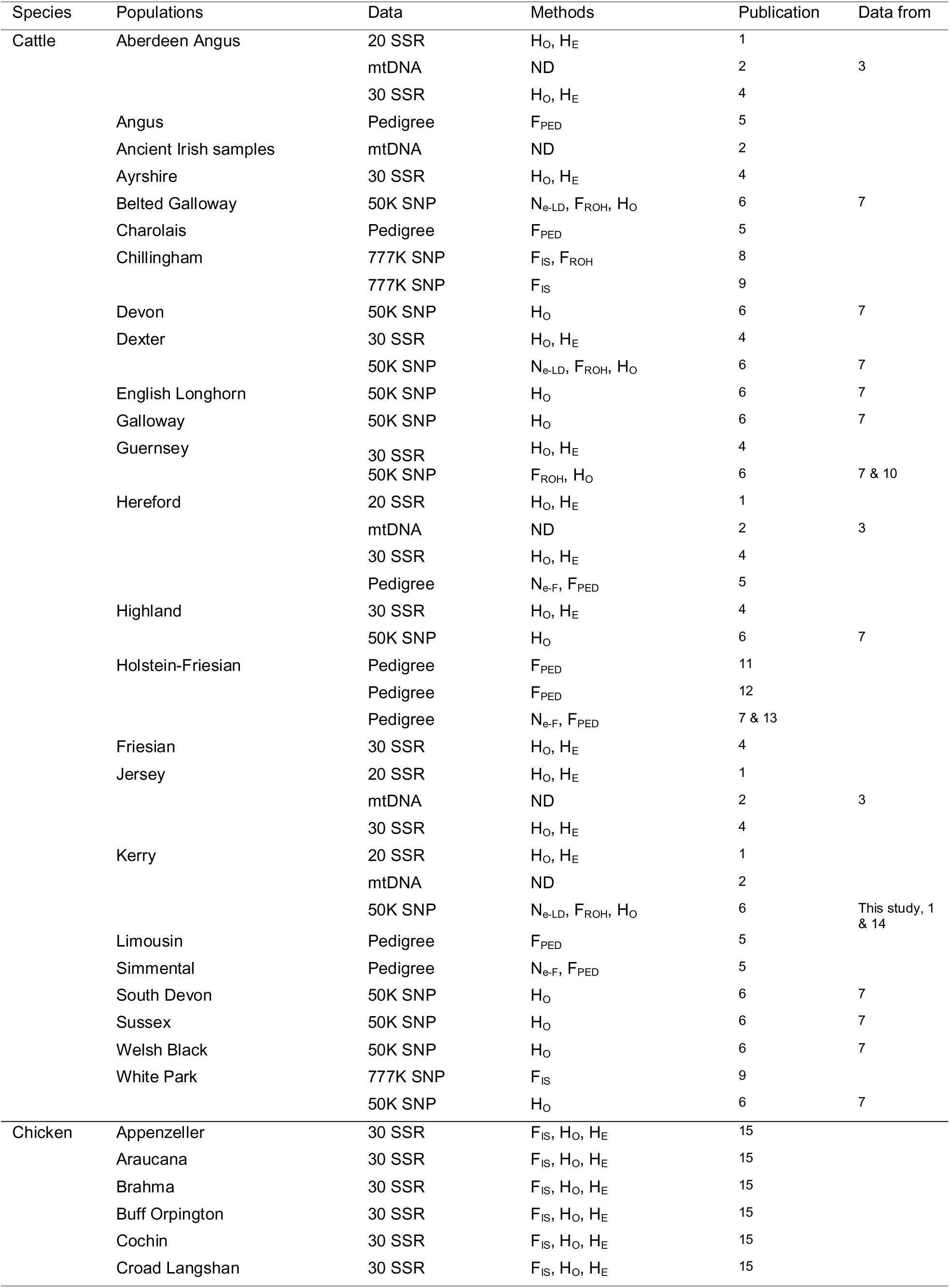
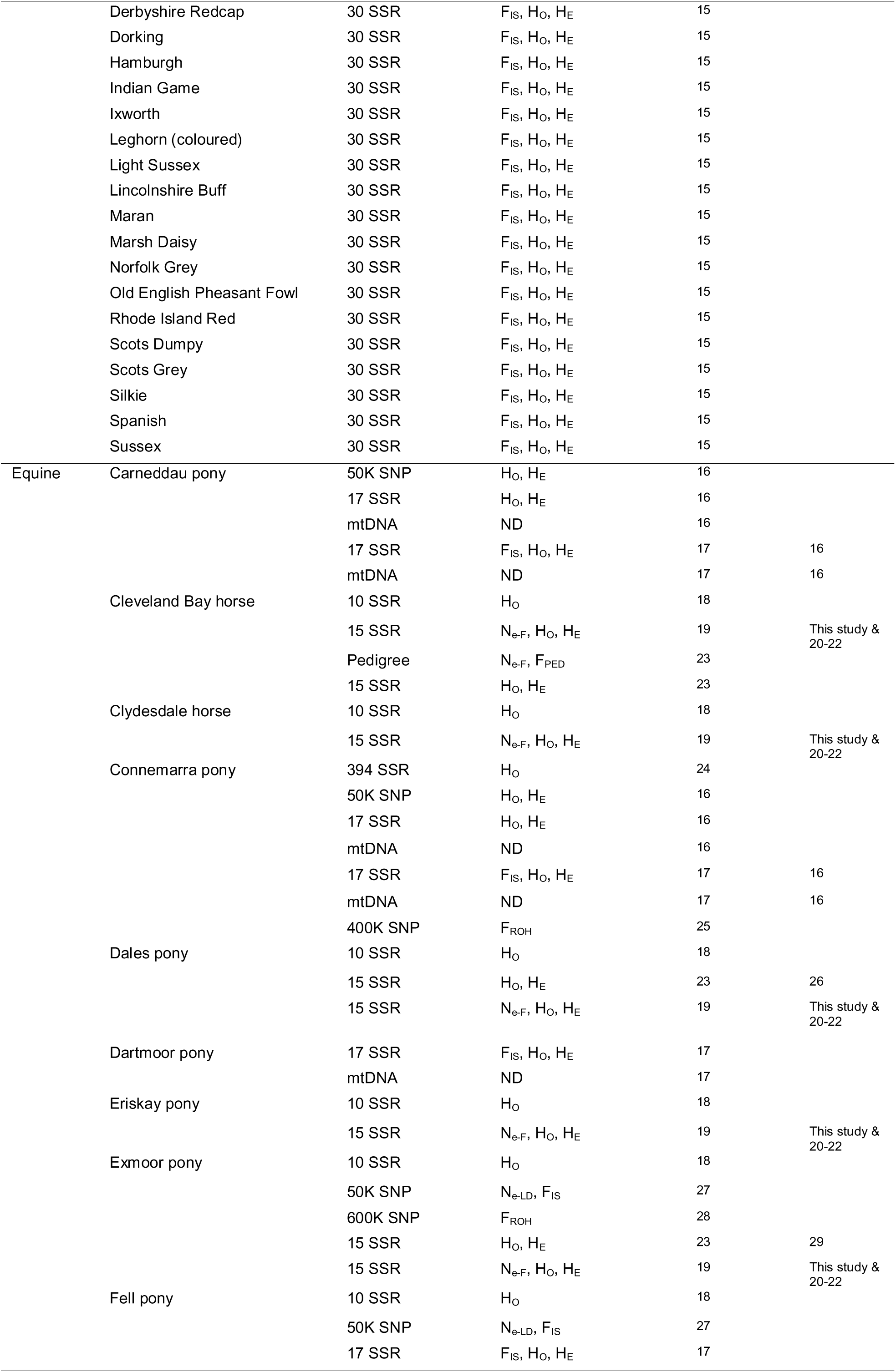
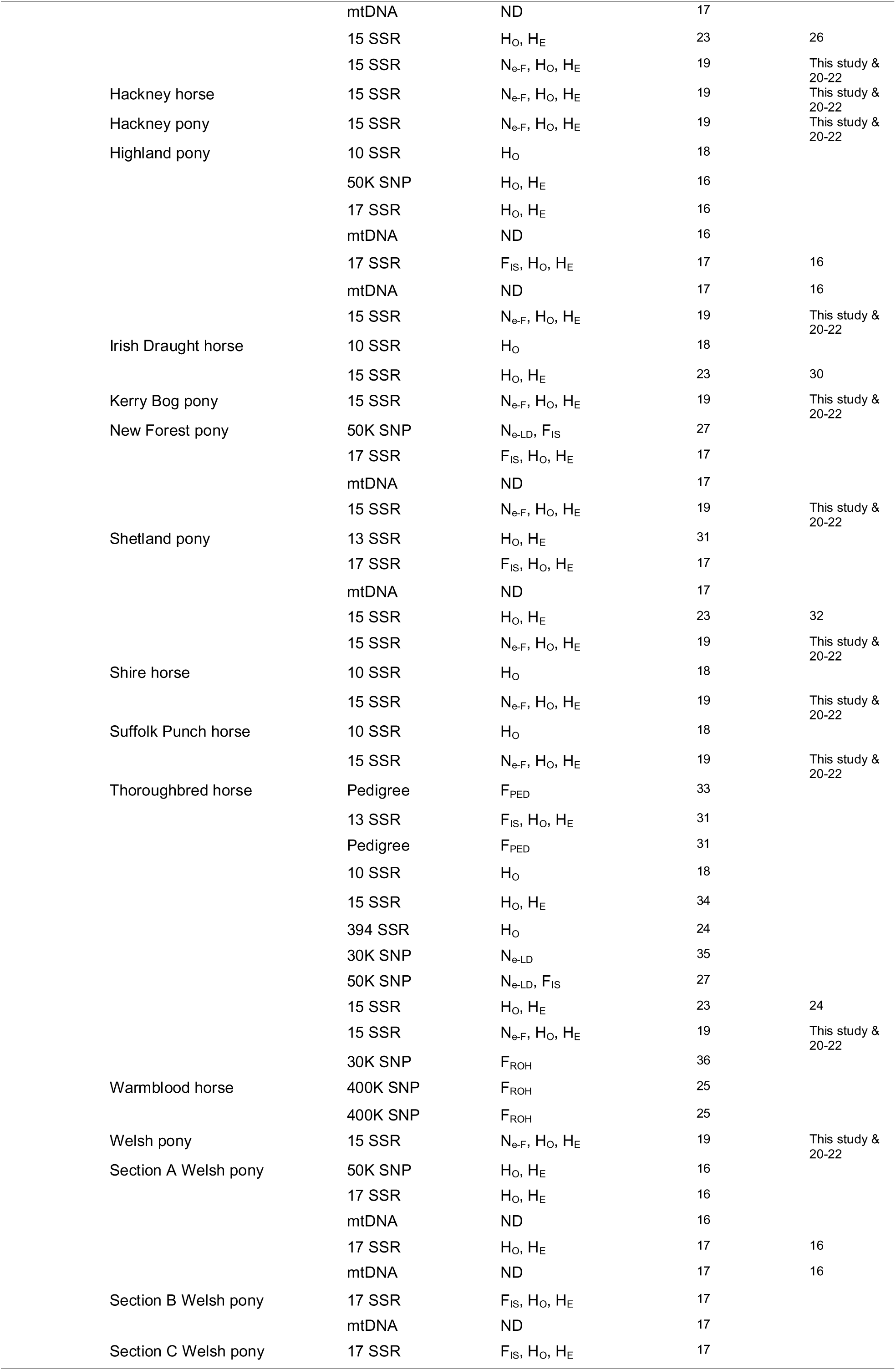
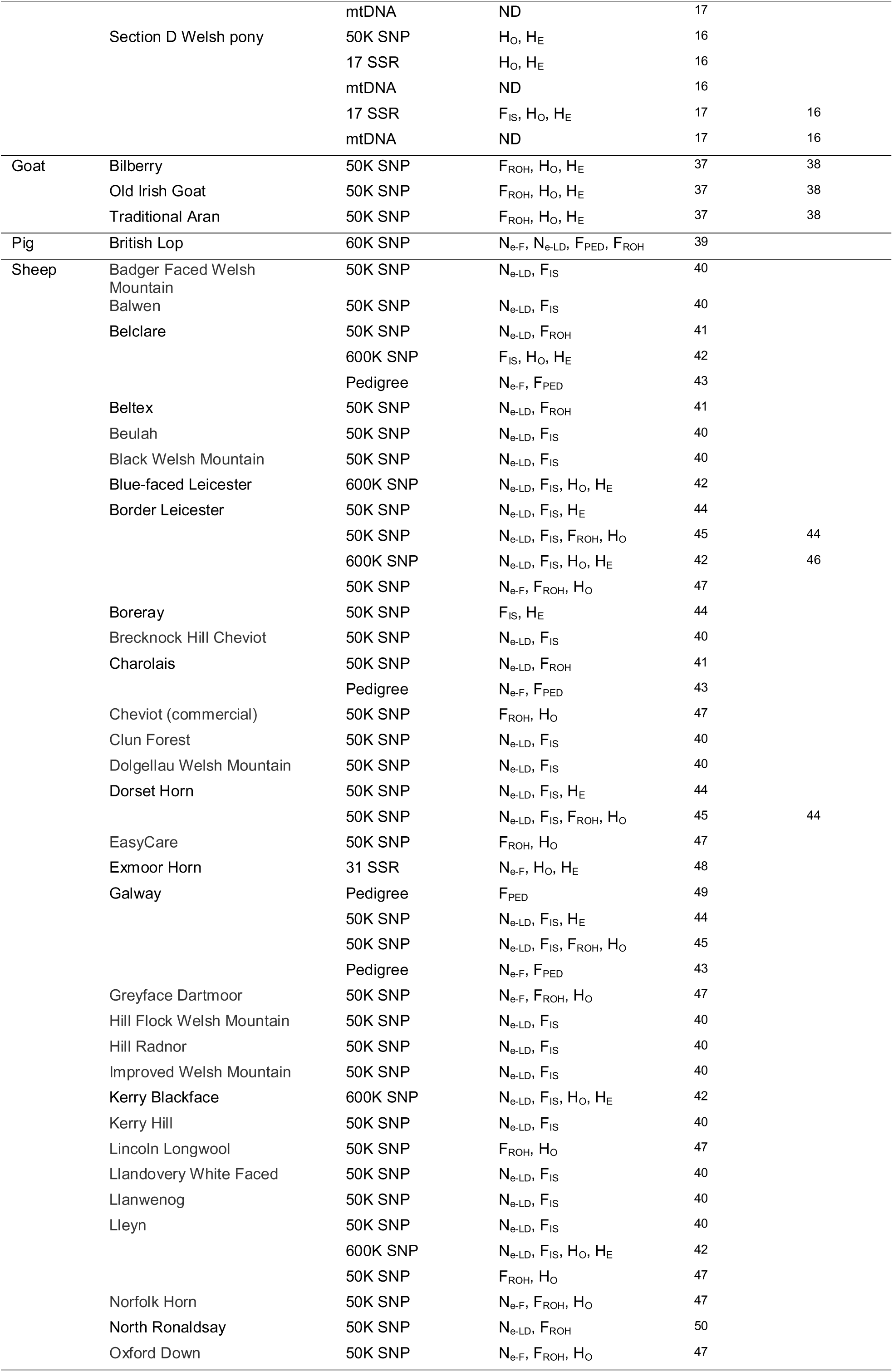
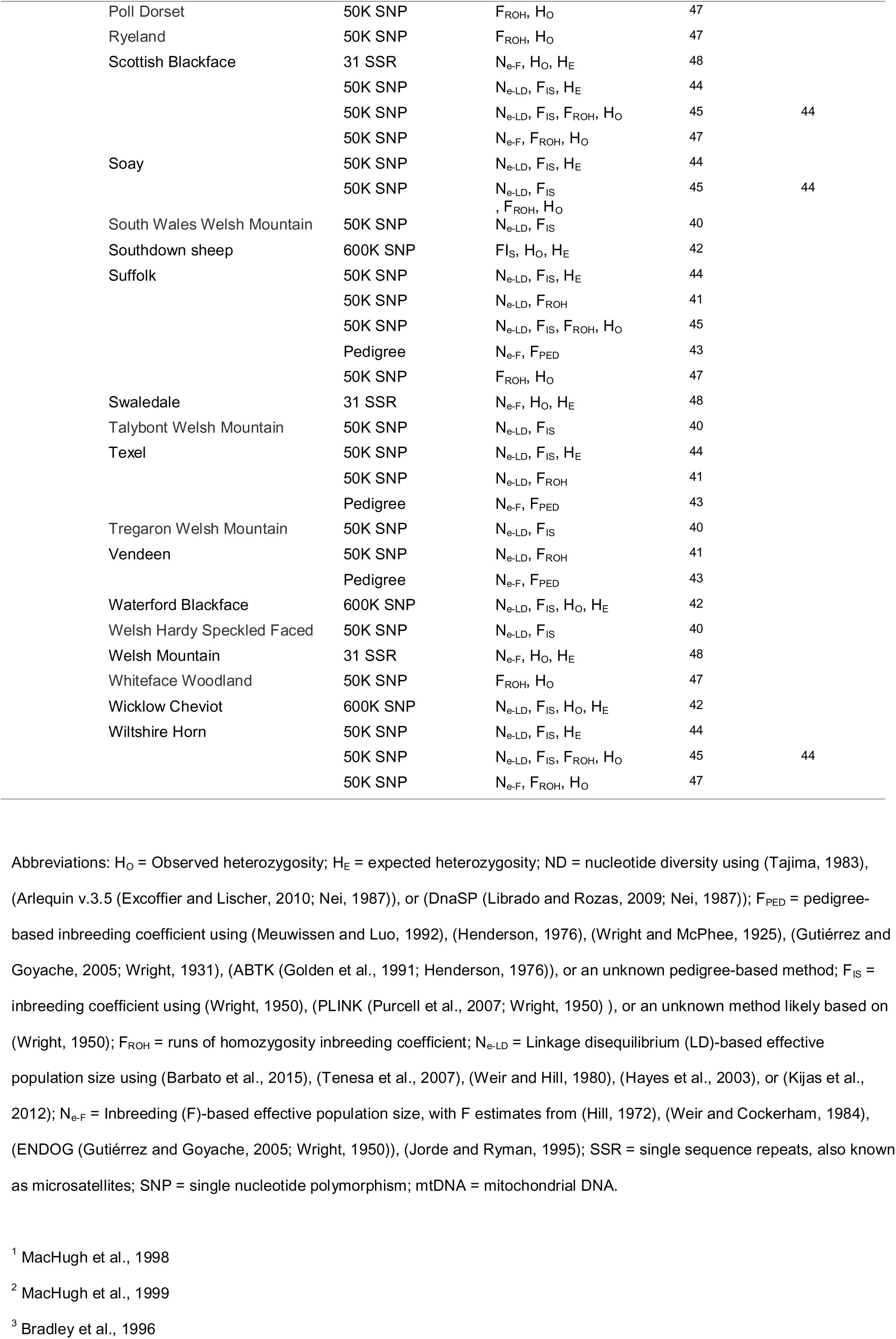
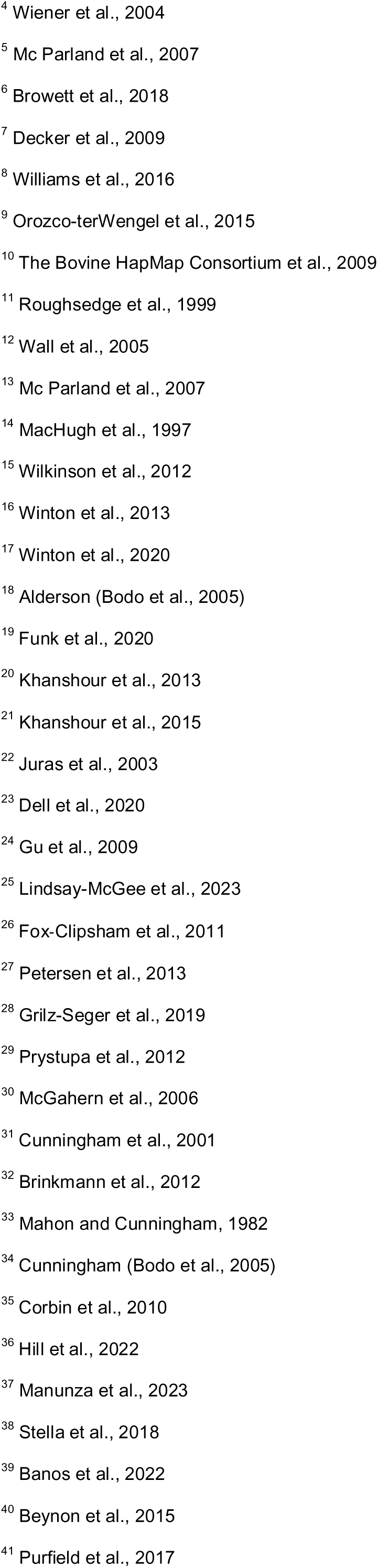
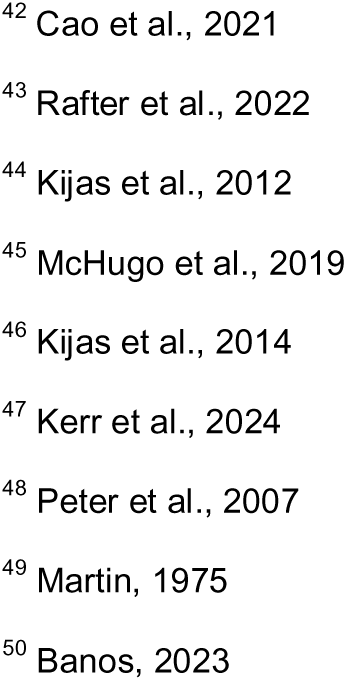
Publications characterising genetic diversity of United Kingdom and/or Irish livestock and equine populations.

### Summary of counts

Of the publications included in our analyses, there were 14 papers on equine populations, 11 on cattle, ten on sheep, one on pig, one on goat and one on chicken. Information on breeds studied, and data type are included in **Table 1**. The number of populations in both the literature and the Rare Breeds Survival Trust 2024 watchlist are plotted by conservation priority status in **Figure 2**. Notably, there were no goat, turkey, duck, or geese populations on the Rare Breeds Survival Trust 2024 watchlist that have estimates of genetic inbreeding, diversity or N_e_ in the literature.

**Figure 2:**
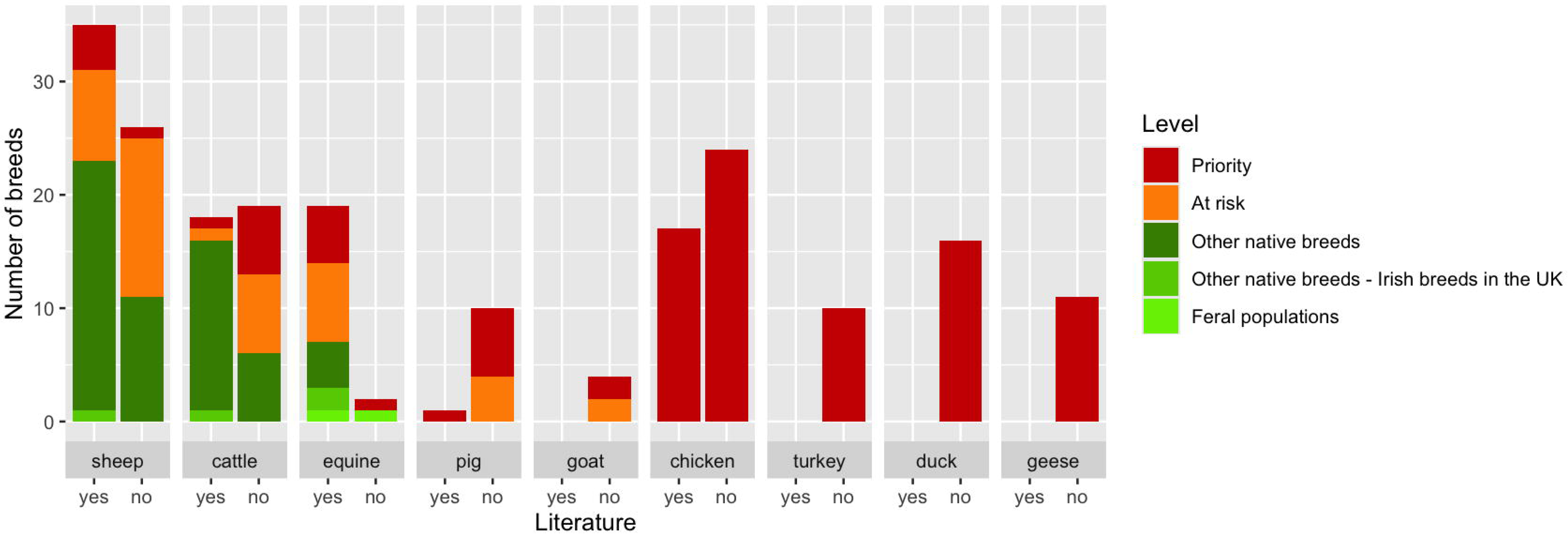
Clustered stacked bar plot of the number of United Kingdom and Irish livestock populations on the Rare Breeds Survival Trust’s watchlist with estimates of inbreeding, diversity or effective population size estimated using pedigree or molecular markers available (“yes”) or not (“no”).

### Comparing genomic-based diversity estimates with census data

Of the N_e_ estimates from genomic data with census estimates available, 43 (of 53) census estimates were greater than inbreeding- or linkage disequilibrium (**LD**)-based estimates. The paired t-test for census-based and inbreeding or LD-based N_e_ was significant (p=0.0009). The median difference between the two was 391 (maximum difference was 24,068). All census-based N_e_ estimates with their corresponding inbreeding- or LD-based N_e_ estimates from the Department for Environment, Food and Rural Affairs are plotted in **Figure 3**.

**Figure 3:**
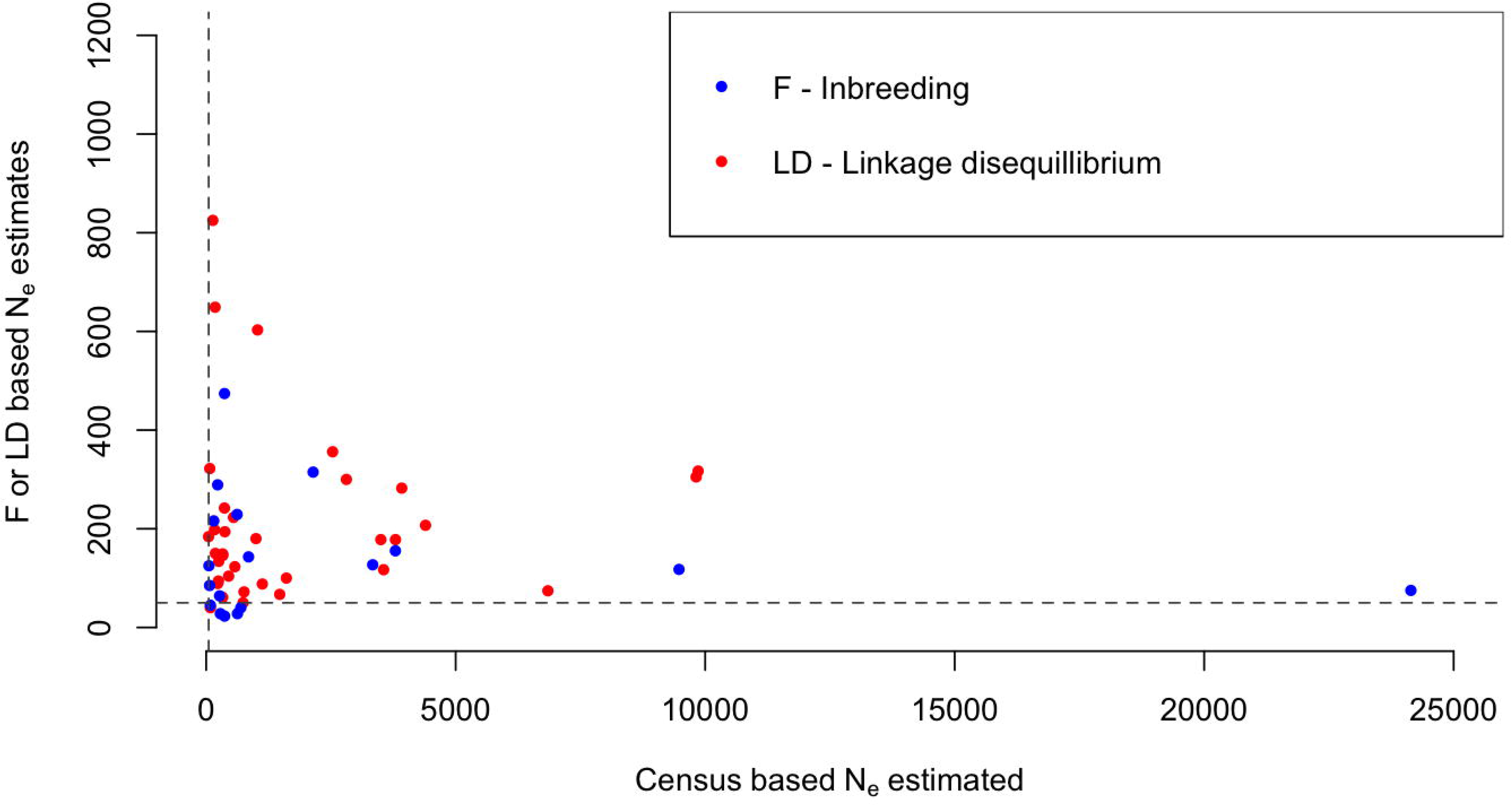
Inbreeding (F)- and linkage disequilibrium (LD)-based effective population size (N_e_) estimates of cattle, equine, pig, and sheep breeds. N_e_ of 50 are indicated on both axes with grey dashed lines.

## Discussion

It is important to maintain rare breeds to adapt to future challenges, to serve niche markets, for conservation grazing, and to maintain cultural identity and contribute to sustainable agriculture. However, there are finite resources available for conservation of these breeds, and they are not immediately profitable in the larger economy, so it is important to identify gaps, and prioritise certain aspects of livestock conservation. In this sense, we discuss: (1) work already done in these areas and compare census-based N_e_ estimates with DNA-based estimates, (2) potential value of whole genome sequence data for rare breeds, (3) usage of N_e_ estimates to guide conservation strategies, (4) limitations of our study design and (5) future actions.

### Effective population size estimates

The well-established threshold of concern for population conservation is a N_e_ of 50 (FAO Commission on Genetic Resources for Food and Agriculture, 2015). Franklin (1980) first proposed a short-term aim of a N_e_ of 50 for small populations (with livestock populations particularly in mind) as this corresponds to a 1% rate of inbreeding (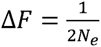) (Wright, 1931), which is considered a tolerable level of inbreeding, considering the negative effect of inbreeding depression. This was part of the 50/500 rule, where Franklin also proposed a long-term minimum value for N_e_ of 500 (where mutation offsets the loss of genetic variation through drift) (Franklin, 1980). There have been more recent discussions on updating the 50/500 rule, which is needed as our knowledge on genome biology has evolved in the following 45 years, however there has been no clear consensus on changing the recommended minimum N_e_ (Waples, 2025; Pérez-Pereira et al., 2022; Frankham et al., 2014; Jamieson and Allendorf, 2012).

The N_e_ threshold of concern may need to be updated, but N_e_ estimates as a measure of genetic diversity also needs to be revisited. We compared census-based and inbreeding- or LD-based N_e_ estimates using annually reported census-based estimates from the Department for Environment, Food and Rural Affairs and inbreeding- or LD-based estimates from the literature. Census-based N_e_ estimates were, on average, higher than inbreeding- or LD-based estimates (with a median difference of 391), and ∼11% (6 of 53) of paired estimates had an inbreeding or LD- based N_e_ estimate < 50, with a corresponding census-based N_e_ > 50. This could indicate that several populations without LD-based N_e_ estimates but with census-based estimates appear to have an N_e_ > 50 but may actually need conservation prioritisation.

We have observed very different N_e_ estimates based on data and methodology. Estimating and publishing census-based estimates has been valuable as a low-cost method for estimating N_e_ and observing trends over time. However, inbreeding and LD-based methods are estimates based on the genome, the diversity of which we are trying to conserve. The difference between estimates could be expected due to the assumptions of the methods used. In the census-based methods, there is an assumption of an idealised population, which does not reflect the reality of livestock breeding. LD-based methods for estimating N_e_ also have assumptions, for example, an assumption is that LD is produced solely from the finite population size, and non-random mating and population structure are absent (Waples, 2025). Additionally, any method using microsatellite or SNP arrays will also contend with ascertainment bias of those DNA markers, where those markers have been identified usually using animals not from the population of interest – which is especially problematic for monitoring rare breeds with unique variants that we are interested in conserving.

The census-based N_e_ estimates in this study showed that more common breeds in the United Kingdom had higher estimates. An extreme example of this is the Holstein-Friesian breed. The United Kingdom census-based N_e_ estimate for 2023 was 13 154 (Beale, 2024). Holstein-Friesian is a globally popular commercial breed, and populations in different countries are well-connected to each other, particularly through similar genetics (global use of artificial insemination with elite bulls’ semen). Therefore, we would expect N_e_ estimates across countries to be somewhat similar to the United Kingdom estimate. N_e_ estimates from the Netherlands (one of the countries where this breed originated from), Canada, France, Ireland, Italy, and the United States of America range from 18 to 150 (Ablondi et al., 2022; Makanjuola et al., 2020; Harmen P. Doekes et al., 2018; H.P. Doekes et al., 2018; Rodríguez-Ramilo et al., 2015; Danchin-Burge et al., 2011; Mc Parland et al., 2007a; Weigel, 2001). These estimates came from a variety of inbreeding- and LD-based methods (but not the census-based method). This highlights how estimating and comparing N_e_estimates can be challenging (even the difference between 18 and 150 has vastly different implications for conservation), and that the census-based United Kingdom estimate is unrealistic. The footnotes of Table 1 demonstrate the wide range of diversity estimates, and even methodologies within diversity estimate type, all with certain assumptions and strengths, rendering meaningful comparisions and intrepretation of results challenging. In the case of census-based N_e_ estimates for livestock populations there is often higher selection intensity in males (fewer active breeding males than active breeding females), which can be accounted for in

Wright’s estimate (Wright, 1931). However, the definition of “active breeding males” could contribute to unreasonable N_e_ estimates. For example, some of the active breeding males (e.g., elite bulls) likely have thousands of offspring, while many active males only have a very small number of offspring, and this variation in family size is not accounted for in the census-based estimate.

### Whole-genome sequencing

While genotyping strategies are becoming more affordable, especially SNP arrays, typically rare breeds are raised as a hobby, and it is not financially feasible to have continual producer-investment in routine genotyping. For producers raising rare breeds for conservation purposes, SNP arrays are not ideal because they likely will not contain unique breed-specific variants (due to ascertainment bias, as seen in Geibel et al. (2021)), which are the main interest of conservation programmes. Next generation sequencing technologies would be better for conservation studies, however, they can be more expensive. Whole genome sequencing costs are decreasing, and there are strategies to reduce those costs even further, including using low-pass sequencing and imputation, but this requires an initial investment to establish high coverage genotyping of a reference population to identify breed-unique variants.

Only one publication included in this study used 15X coverage Illumina 150 base pair paired-end whole genome sequence data (Lindsay-McGee et al., 2023). Due to the study design which also used available genotypes from populations genotyped with an Affymetrix array, only 400K SNPs remained for analyses. In contrast, to give an idea of the magnitude of the number of variants we could expect from sequencing, from a publication in 2019 from the 1000 Bull Genome Project where a total of 429 animals from 14 breeds were sequenced, they identified 29.1 million single nucleotide variants, and 1.7 million insertions-deletions (Hayes et al., 2014). We could expect that there is a lot of unexplored data from the Lindsay-McGee et al. (2023) study, and that it would be valuable for further investment in generating whole genome sequence data for other rare breeds in the United Kingdom, allowing for information on breed-unique variants to be included in management plans.

### Conservation management

As we previously mentioned, monitoring and follow-up conservation strategies are mainly based on the Food and Agriculture Organisation’s guidance (N_e_ of 50 per generation, equivalent to 1% inbreeding rate per generation (FAO Commission on Genetic Resources for Food and Agriculture, 2015)). Current census-based estimates of N_e_ should be replaced by inbreeding-based estimates of N_e_. For the census-based estimates of N_e_, the Department for Environment, Food and Rural Affairs is using the number of active sires and dams, which is known because pedigree information is collected and available. An immediate improvement would be to use pedigree information to estimate rates of inbreeding, which could then be used to calculate annual inbreeding-based N_e_ estimates. In the long-term, developing a national strategy to regularly collect DNA samples for breeds on the Rare Breeds Survival Trust watchlist, would allow for DNA-based estimates of genetic diversity measures.

Having DNA-based contemporary N_e_ estimates is critical to prioritise breeds/populations for conservation activities. Furthermore, having genomic (especially whole genome sequence) data on these breeds would not only allow for better N_e_ estimates but would also lead to a better understanding of the general population structure. This would add value for cost-effective decision making, particularly linked to choosing the most optimal conservation system (*in situ*, *ex situ in vivo*, *ex situ in vitro*), optimising the number of individuals within the breed to keep in the chosen conservation system, as well as opening the door for future activities (e.g., precise introgression, genome editing, etc.). Conservation activities and systems based on genomic data were systematically reviewed in Oldenbroek et al. (2017), the algorithms for selection and optimisation of the number of individuals are available and can be easily adapted for conservation purposes, e.g., Pocrnic et al., (2022); Yu et al., (2014), and for a recent review of the genome editing concepts applied to livestock populations, including conservation applications, see Clark, (2020). We particularly mention genome editing technology as there is potential to introduce breed-unique causal variants into other populations (Leroy et al., 2016).

Alternatively, variants could be introduced by genome editing to make local populations more resistant to disease or improve their fertility (traits that may suffer from inbreeding depression), only changing traits to improve the survival of the breed while maintaining the breed standards/definitions. Finally, a more complex application of genome editing that could be considered is surrogate sire technology. Surrogate sires are made sterile by CRISPR-Cas9 editing of *NANOS2* in mammals (Park et al., 2017) or *DDX4* in chicken (Taylor et al., 2017), and donor-derived stem cells are transplanted into the gonads (Taylor et al., 2017). The surrogate sire could then disseminate the donor’s sperm through natural breeding. The potential to exploit surrogate sire technology in commercial livestock breeding was discussed by Gottardo et al. (2019), but similarly, we could use it in a conservation context to disseminate local breed sperm. Finally, novel technologies like gene drives are emerging and could find their application in conservation strategies to help spread desired alleles in a population (e.g., Faber et al. (2021); Webber et al. (2015); Esvelt et al. (2014)).

### Limitations of this study

Many of the results included in this study either came from or benefitted from international research consortiums. These consortiums build capacity through the training of individuals, building databases and sharing resources. However, from these consortium publications there were several breeds that originated in the United Kingdom, that were not included in the current study because of uncertainty of sampling locations. Additionally, breeds originating in the United Kingdom with populations in other countries, may no longer reflect the current United Kingdom population anymore due to the use of crossbreeding. Therefore, we did not include populations unless we were certain that they were United Kingdom or Ireland samples and if there was a proportion of samples included from another country, there was some evidence that these populations were well-connected with the corresponding population in the United Kingdom.

There are more peer-reviewed publications that analyse genotypes from livestock populations in the United Kingdom than we have included in this publication. We tried to keep only those publications with some genetic diversity measure, inbreeding or N_e_ estimate. This means papers looking at population structure and signatures of selections, while related, were not included. Sometimes the data from those studies were used in other publications where the measures we were interested in were estimated, and we tried to cite both the original source of data and the publication with the analyses of interest.

From systematically reviewing literature for genetic diversity, inbreeding and N_e_ estimates for livestock and equine populations in the United Kingdom and Ireland, one recommendation for other researchers is clear: include detailed information on populations analysed, including sampling year, number of herds sampled from and sampling criteria. This additional information will be helpful for future studies (for publicly available data eventually used in other studies) and can be used to inform conservation decisions.

### Future actions

This study was designed to identify gaps in our knowledge of the genetic diversity of livestock populations in the United Kingdom. Most breeds on the Rare Breeds Survival Trust watchlist do not have DNA-based studies, and for those with estimates, many are outdated. Most notably missing were any estimates on duck, geese, goat and turkey breeds. These populations should be prioritised for generating genomic data, and they need to remain on the Rare Breeds Survival Trust watchlist and the Department for Environment, Food and Rural Affairs monitoring programmes, potentially even reassessing conservation categories based on DNA-based metrics when available. The creation of a centralised database for all rare livestock breeds in the United Kingdom, integrating the Department for Environment, Food and Rural Affairs, Rare Breeds Survival Trust and academic sources, would help with implementing a systematic approach to monitor genetic diversity, and fund the generation of genomic data to update diversity estimates. Publicly funded genetic estimates and resulting publications should be open access and allow for data sharing. Through the standardisation and sharing of data, joint conservation programmes for transboundary breeds can be developed, and breed connectedness and genetic diversity measures can be jointly assessed.

## Ethics approval

Ethics approval was not needed for this study.

## Data and model availability statement

The data were not deposited in an official repository. All data are available in the supplemental information.

## Declaration of generative AI and AI-assisted technologies in the writing process

We declare that no generative AI or AI-assisted technologies were used in the writing process.

## Acknowledgements

For the purpose of open access, the authors have applied a CC BY public copyright license to any author accepted manuscript version arising from this submission. This article was deposited as a pre-print with bioRxiv (doi: https://doi.org/10.1101/2025.05.14.653764).

## Supplementary files

**Supplementary Table S1:** Supplementary_file_1.xlsx – Effective population size, inbreeding, observed and expected heterozygosity, and nucleotide diversity of United Kingdom and Irish livestock and equine populations

## References

Ablondi, M., Sabbioni, A., Stocco, G., Cipolat-Gotet, C., Dadousis, C., Kaam, J.-T.V., Finocchiaro, R., Summer, A., 2022. Genetic Diversity in the Italian Holstein Dairy Cattle Based on Pedigree and SNP Data Prior and After Genomic Selection. Frontiers in Veterinary Science 8, 773985. doi:10.3389/fvets.2021.773985

Banos, G., 2023. Genomic analysis of the North Ronaldsay sheep. Rare Breeds Survival Trust (RBST), Kenilworth, United Kingdom. Retrieved on 8 September 2025 from https://www.rbst.org.uk/Handlers/Download.ashx?IDMF=d71f4977-b219-479f-be8b-b70702988263

Banos, G., Talenti, A., Chatziplis, D., Sánchez-Molano, E., 2022. Genomic analysis of the rare British Lop pig and identification of distinctive genomic markers. PLOS ONE 17, e0271053. doi:10.1371/journal.pone.0271053

Barbato, M., Orozco-Terwengel, P., Tapio, M., Bruford, M.W., 2015. SNeP: a tool to estimate trends in recent effective population size trajectories using genome-wide SNP data. Frontiers in Genetics 6, 109. doi:10.3389/fgene.2015.00109

Beale, S., 2024. Official Statistics UK farm animal genetic resources (FAnGR): breed inventory results. UK Department for Environment, Food & Rural Affairs. Retrieved on 8 September 2025 from https://www.gov.uk/government/statistics/uk-farm-animal-genetic-resources-fangr-breed-inventory-results

Beynon, S.E., Slavov, G.T., Farré, M., Sunduimijid, B., Waddams, K., Davies, B., Haresign, W., Kijas, J., MacLeod, I.M., Newbold, C.J., Davies, L., Larkin, D.M., 2015. Population structure and history of the Welsh sheep breeds determined by whole genome genotyping. BMC Genetics 16, 65. doi:10.1186/s12863-015-0216-x

Bodo, I., Alderson, L., Langlois, B. (Eds), 2005. Conservation genetics of endangered horse breeds. EAAP Scientific Series, Wageningen Academic Publishers, Wageningen, Netherlands. Retrieved on 8 September 2025 from https://brill.com/edcollbook/title/68563?language=en&srsltid=AfmBOorAnLyai-phwpa_e2vwhxyGbIRGn70_5iUyppsgFkkea8d1VgKS

Bradley, D.G., MacHugh, D.E., Cunningham, P., Loftus, R.T., 1996. Mitochondrial diversity and the origins of African and European cattle. Proceedings of the National Academy of Sciences 93, 5131–5135. doi:10.1073/pnas.93.10.5131

Brinkmann, L., Gerken, M., Riek, A., 2012. Adaptation strategies to seasonal changes in environmental conditions of a domesticated horse breed, the Shetland pony (*Equus ferus caballus*). Journal of Experimental Biology 215, 1061–1068. doi:10.1242/jeb.064832

Browett, S., McHugo, G., Richardson, I.W., Magee, D.A., Park, S.D.E., Fahey, A.G., Kearney, J.F., Correia, C.N., Randhawa, I.A.S., MacHugh, D.E., 2018. Genomic Characterisation of the Indigenous Irish Kerry Cattle Breed. Frontiers in Genetics 9, 51. doi:10.3389/fgene.2018.00051

Canali, G., 2006. Common agricultural policy reform and its effects on sheep and goat market and rare breeds conservation. Small Ruminant Research 62, 207–213. doi:10.1016/j.smallrumres.2005.08.021

Cao, Y.-H., Xu, S.-S., Shen, M., Chen, Z.-H., Gao, L., Lv, F.-H., Xie, X.-L., Wang, X.-H., Yang, H., Liu, C.-B., Zhou, P., Wan, P.-C., Zhang, Y.-S., Yang, J.-Q., Pi, W.-H., Hehua, Ee., Berry, D.P., Barbato, M., Esmailizadeh, A., Nosrati, M., Salehian-Dehkordi, H., Dehghani-Qanatqestani, M., Dotsev, A.V., Deniskova, T.E., Zinovieva, N.A., Brem, G., Štěpánek, O., Ciani, E., Weimann, C., Erhardt, G., Mwacharo, J.M., Ahbara, A., Han, J.-L., Hanotte, O., Miller, J.M., Sim, Z., Coltman, D., Kantanen, J., Bruford, M.W., Lenstra, J.A., Kijas, J., Li, M.-H., 2021. Historical Introgression from Wild Relatives Enhanced Climatic Adaptation and Resistance to Pneumonia in Sheep. Molecular Biology and Evolution 38, 838–855. doi:10.1093/molbev/msaa236

Clark, E.L., 2020. Breeding in an Era of Genome Editing. In Encyclopedia of Sustainability Science and Technology (ed. Meyers, R.A.). Springer New York, New York, NY, pp. 1–16. doi:10.1007/978-1-4939-2493-6_1122-1

Corbin, L.J., Blott, S.C., Swinburne, J.E., Vaudin, M., Bishop, S.C., Woolliams, J.A., 2010. Linkage disequilibrium and historical effective population size in the Thoroughbred horse. Animal Genetics 41, 8–15. doi:10.1111/j.1365-2052.2010.02092.x

Cunningham, E.P., Dooley, J.J., Splan, R.K., Bradley, D.G., 2001. Microsatellite diversity, pedigree relatedness and the contributions of founder lineages to thoroughbred horses. Animal Genetics 32, 360–364. doi:10.1046/j.1365-2052.2001.00785.x

Danchin-Burge, C., Hiemstra, S.J., Blackburn, H., 2011. Ex situ conservation of Holstein-Friesian cattle: Comparing the Dutch, French, and US germplasm collections. Journal of Dairy Science 94, 4100–4108. doi:10.3168/jds.2010-3957

Decker, J.E., Pires, J.C., Conant, G.C., McKay, S.D., Heaton, M.P., Chen, K., Cooper, A., Vilkki, J., Seabury, C.M., Caetano, A.R., Johnson, G.S., Brenneman, R.A., Hanotte, O., Eggert, L.S., Wiener, P., Kim, J.-J., Kim, K.S., Sonstegard, T.S., Van Tassell, C.P., Neibergs, H.L., McEwan, J.C., Brauning, R., Coutinho, L.L., Babar, M.E., Wilson, G.A., McClure, M.C., Rolf, M.M., Kim, J., Schnabel, R.D., Taylor, J.F., 2009. Resolving the evolution of extant and extinct ruminants with high-throughput phylogenomics. Proceedings of the National Academy of Sciences 106, 18644–18649. doi:10.1073/pnas.0904691106

DEFRA, 2021. Policy paper: Native livestock breeds: reducing extinction risk. Department for Environment, Food & Rural Affairs. Retrieved on 8 September 2025 from https://www.gov.uk/government/publications/native-livestock-breeds-reducing-extinction-risk/native-livestock-breeds-reducing-extinction-risk

Dell, A., Curry, M., Yarnell, K., Starbuck, G., Wilson, P.B., 2020. Genetic analysis of the endangered Cleveland Bay horse: A century of breeding characterised by pedigree and microsatellite data. PLOS ONE 15, e0240410. doi:10.1371/journal.pone.0240410

Doekes, H.P., Veerkamp, R.F., Bijma, P., Hiemstra, S.J., Windig, J., 2018. Value of the Dutch Holstein Friesian germplasm collection to increase genetic variability and improve genetic merit. Journal of Dairy Science 101, 10022– 10033. doi:10.3168/jds.2018-15217

Doekes, Harmen P., Veerkamp, R.F., Bijma, P., Hiemstra, S.J., Windig, J.J., 2018. Trends in genome-wide and region-specific genetic diversity in the Dutch-Flemish Holstein–Friesian breeding program from 1986 to 2015. Genetics Selection Evolution 50, 15. doi:10.1186/s12711-018-0385-y

Esvelt, K.M., Smidler, A.L., Catteruccia, F., Church, G.M., 2014. Concerning RNA-guided gene drives for the alteration of wild populations. eLife 3, e03401. doi:10.7554/eLife.03401

Excoffier, L., Lischer, H.E.L., 2010. Arlequin suite ver 3.5: a new series of programs to perform population genetics analyses under Linux and Windows. Molecular Ecology Resources 10, 564–567. doi:10.1111/j.1755-0998.2010.02847.x

Faber, N.R., McFarlane, G.R., Gaynor, R.C., Pocrnic, I., Whitelaw, C.B.A., Gorjanc, G., 2021. Novel combination of CRISPR-based gene drives eliminates resistance and localises spread. Scientific Reports 11, 3719. doi:10.1038/s41598-021-83239-4

Falconer, D.S., Mackay, T.F.C., 1996. Introduction to Quantitative Genetics. Pearson Education Limited, Harlow, England.

FAO Commission on Genetic Resources for Food and Agriculture, 2015. The second report on the state of the world’s animal genetic resorces for food and agriculture. FAO, Rome, Italy.

Fox_-_Clipsham, L.Y., Brown, E.E., Carter, S.D., Swinburne, J.E., 2011. Population screening of endangered horse breeds for the foal immunodeficiency syndrome mutation. Veterinary Record 169, 655–655. doi:10.1136/vr.100235

Frankham, R., Bradshaw, C.J.A., Brook, B.W., 2014. Genetics in conservation management: Revised recommendations for the 50/500 rules, Red List criteria and population viability analyses. Biological Conservation 170, 56–63. doi:10.1016/j.biocon.2013.12.036

Franklin, 1980. Evolutionary changes in small populations. In Conservation Biology. An Evolutionary-Ecological Perspective. Sinauer Associates, Sunderland, MA, USA, pp. 135–149.

Funk, S.M., Guedaoura, S., Juras, R., Raziq, A., Landolsi, F., Luís, C., Martínez, A.M., Musa Mayaki, A., Mujica, F., Oom, M.D.M., Ouragh, L., Stranger, Y., Vega_-_Pla, J.L., Cothran, E.G., 2020. Major inconsistencies of inferred population genetic structure estimated in a large set of domestic horse breeds using microsatellites. Ecology and Evolution 10, 4261–4279. doi:10.1002/ece3.6195

Geibel, J., Reimer, C., Weigend, S., Weigend, A., Pook, T., Simianer, H., 2021. How array design creates SNP ascertainment bias. PLOS ONE 16, e0245178. doi:10.1371/journal.pone.0245178

Geographical indications and quality schemes explained, n.d. . European Commission. Retrieved on 8 September 2025 from https://agriculture.ec.europa.eu/farming/geographical-indications-and-quality-schemes/geographical-indications-and-quality-schemes-explained_en

Golden, B.L., Brinks, J.S., Bourdon, R.M., 1991. A performance programmed method for computing inbreeding coefficients from large data sets for use in mixed-model analyses. Journal of Animal Science 69, 3564–3573. doi:10.2527/1991.6993564x

Gottardo, P., Gorjanc, G., Battagin, M., Gaynor, R.C., Jenko, J., Ros-Freixedes, R., Bruce A. Whitelaw, C., Mileham, A.J., Herring, W.O., Hickey, J.M., 2019. A Strategy To Exploit Surrogate Sire Technology in Livestock Breeding Programs. G3 Genes|Genomes|Genetics 9, 203–215. doi:10.1534/g3.118.200890

Graze with livestock to maintain and improve habitats, n.d.. UK Department for Environment, Food & Rural Affairs. Retrieved on 8 September 2025 from https://defrafarming.blog.gov.uk/graze-with-livestock-to-maintain-and-improve-habitats/

Grilz-Seger, G., Neuditschko, M., Ricard, A., Velie, B., Lindgren, G., Mesarič, M., Cotman, M., Horna, M., Dobretsberger, M., Brem, G., Druml, T., 2019. Genome-Wide Homozygosity Patterns and Evidence for Selection in a Set of European and Near Eastern Horse Breeds. Genes 10, 491. doi:10.3390/genes10070491

Gu, J., Orr, N., Park, S.D., Katz, L.M., Sulimova, G., MacHugh, D.E., Hill, E.W., 2009. A Genome Scan for Positive Selection in Thoroughbred Horses. PLoS ONE 4, e5767. doi:10.1371/journal.pone.0005767

Gutiérrez, J.P., Goyache, F., 2005. A note on ENDOG: a computer program for analysing pedigree information. Journal of Animal Breeding and Genetics 122, 172–176. doi:10.1111/j.1439-0388.2005.00512.x

Hayes, B.J., Macleod, I.M., Daetwyler, H.D., Bowman, P.J., Chamberlian, A.J., Jagt, C.J.V., Capitan, A., Pausch, H., Stothard, P., Liao, X., Schrooten, C., Mullaart, E., Fries, R., Guldbrandtsen, B., Lund, M.S., Boichard, D., Veerkamp, R.F., Vantassell, C.P., Gredler, B., Druet, T., Bagnato, A., Vilkki, J., Dekoning, D.J., Santus, E., Goddard, M.E., 2014. Genomic prediction from whole genome sequence in livestock: the 1000 Bull Genomes Project. In Proceedings of the 12th World Congress on Genetics Applied to Livestock Production (WCGALP), 17-22 August 2014, Vancouver, Canada. Retrieved on 8 September 2025 from https://hal.science/hal-01193911v1/document

Hayes, B.J., Visscher, P.M., McPartlan, H.C., Goddard, M.E., 2003. Novel Multilocus Measure of Linkage Disequilibrium to Estimate Past Effective Population Size. Genome Research 13, 635–643. doi:10.1101/gr.387103

Henderson, C.R., 1976. A Simple Method for Computing the Inverse of a Numerator Relationship Matrix Used in Prediction of Breeding Values. Biometrics 32, 69. doi:10.2307/2529339

Hill, E.W., Stoffel, M.A., McGivney, B.A., MacHugh, D.E., Pemberton, J.M., 2022. Inbreeding depression and the probability of racing in the Thoroughbred horse. Proceedings of the Royal Society B: Biological Sciences 289, 20220487. doi:10.1098/rspb.2022.0487

Hill, W.G., 1972. Effective size of populations with overlapping generations. Theoretical Population Biology 3, 278–289. doi:10.1016/0040-5809(72)90004-4

Jamieson, I.G., Allendorf, F.W., 2012. How does the 50/500 rule apply to MVPs? Trends in Ecology & Evolution 27, 578–584. doi:10.1016/j.tree.2012.07.001

Jorde, P.E., Ryman, N., 1995. Temporal allele frequency change and estimation of effective size in populations with overlapping generations. Genetics 139, 1077–1090. doi:10.1093/genetics/139.2.1077

Juras, R., Cothran, E.G., Klimas, R., 2003. Genetic Analysis of Three Lithuanian Native Horse Breeds. Acta Agriculturae Scandinavica, Section A - Animal Science 53, 180–185. doi:10.1080/09064700310012971

Kerr, E., Marr, M.M., Collins, L., Dubarry, K., Salavati, M., Scinto, A., Woolley, S., Clark, E.L., 2024. Analysis of genotyping data reveals the unique genetic diversity represented by the breeds of sheep native to the United Kingdom. BMC Genomic Data 25, 82. doi:10.1186/s12863-024-01265-3

Khanshour, A., Juras, R., Blackburn, R., Cothran, E.G., 2015. The Legend of the Canadian Horse: Genetic Diversity and Breed Origin. Journal of Heredity 106, 37–44. doi:10.1093/jhered/esu074

Khanshour, A.M., Juras, R., Cothran, E.G., 2013. Microsatellite analysis of genetic variability in Waler horses from Australia. Australian Journal of Zoology 61, 357. doi:10.1071/ZO13062

Kijas, J.W., Lenstra, J.A., Hayes, B., Boitard, S., Porto Neto, L.R., San Cristobal, M., Servin, B., McCulloch, R., Whan, V., Gietzen, K., Paiva, S., Barendse, W., Ciani, E., Raadsma, H., McEwan, J., Dalrymple, B., other members of the International Sheep Genomics Consortium, 2012. Genome-Wide Analysis of the World’s Sheep Breeds Reveals High Levels of Historic Mixture and Strong Recent Selection. PLoS Biology 10, e1001258. doi:10.1371/journal.pbio.1001258

Kijas, J.W., Porto-Neto, L., Dominik, S., Reverter, A., Bunch, R., McCulloch, R., Hayes, B.J., Brauning, R., McEwan, J., the International Sheep Genomics Consortium, 2014. Linkage disequilibrium over short physical distances measured in sheep using a high-density SNP chip. Animal Genetics 45, 754–757. doi:10.1111/age.12197

Larson, G., Piperno, D.R., Allaby, R.G., Purugganan, M.D., Andersson, L., Arroyo-Kalin, M., Barton, L., Climer Vigueira, C., Denham, T., Dobney, K., Doust, A.N., Gepts, P., Gilbert, M.T.P., Gremillion, K.J., Lucas, L., Lukens, L., Marshall, F.B., Olsen, K.M., Pires, J.C., Richerson, P.J., Rubio de Casas, R., Sanjur, O.I., Thomas, M.G., Fuller, D.Q., 2014. Current perspectives and the future of domestication studies. Proceedings of the National Academy of Sciences 111, 6139–6146. doi:10.1073/pnas.1323964111

Leroy, G., Besbes, B., Boettcher, P., Hoffmann, I., Capitan, A., Baumung, R., 2016. Rare phenotypes in domestic animals: unique resources for multiple applications. Animal Genetics 47, 141–153. doi:10.1111/age.12393

Librado, P., Rozas, J., 2009. DnaSP v5: a software for comprehensive analysis of DNA polymorphism data. Bioinformatics 25, 1451–1452. doi:10.1093/bioinformatics/btp187

Lindsay-McGee, V., Sanchez-Molano, E., Banos, G., Clark, E.L., Piercy, R.J., Psifidi, A., 2023. Genetic characterisation of the Connemara pony and the Warmblood horse using a within-breed clustering approach. Genetics Selection Evolution 55, 60. doi:10.1186/s12711-023-00827-w

MacHugh, D.E., Loftus, R.T., Cunningham, P., Bradley, D.G., 1998. Genetic structure of seven European cattle breeds assessed using 20 microsatellite markers. Animal Genetics 29, 333–340. doi:10.1046/j.1365-2052.1998.295330.x

MacHugh, D.E., Shriver, M.D., Loftus, R.T., Cunningham, P., 1997. Microsatellite DNA Variation and the Evolution, Domestication and Phylogeography of Taurine and Zebu Cattle (Bos taurus and Bos indicus). Genetics 146, 1071– 1086. 10.1093/genetics/146.3.1071

MacHugh, D.E., Troy, C.S., McCormick, F., Olsaker, I., Eythórsdóttir, E., Bradle, D.G., 1999. Early medieval cattle remains from a Scandinavian settlement in Dublin: genetic analysis and comparison with extant breeds. Philosophical Transactions of the Royal Society of London. Series B: Biological Sciences 354, 99–109. doi:10.1098/rstb.1999.0363

Mahon, G.A.T., Cunningham, E.P., 1982. Inbreeding and the inheritance of fertility in the thoroughbred mare. Livestock Production Science 9, 743–754. doi:10.1016/0301-6226(82)90021-5

Makanjuola, B.O., Miglior, F., Abdalla, E.A., Maltecca, C., Schenkel, F.S., Baes, C.F., 2020. Effect of genomic selection on rate of inbreeding and coancestry and effective population size of Holstein and Jersey cattle populations. Journal of Dairy Science 103, 5183–5199. doi:10.3168/jds.2019-18013

Manunza, A., Ramirez-Diaz, J., Cozzi, P., Lazzari, B., Tosser-Klopp, G., Servin, B., Johansson, A.M., Grøva, L., Berg, P., Våge, D.I., Stella, A., 2023. Genetic diversity and historical demography of underutilised goat breeds in North-Western Europe. Scientific Reports 13, 20728. doi:10.1038/s41598-023-48005-8

Martin, I., 1975. A Genetic Analysis of the Galway Sheep Breed: 3. Level of Inbreeding in the Pedigree Galway Sheep Breed. Irish Journal of Agricultural Research 14, 269–274.

Mc Parland, S., Kearney, J.F., Rath, M., Berry, D.P., 2007a. Inbreeding Effects on Milk Production, Calving Performance, Fertility, and Conformation in Irish Holstein-Friesians. Journal of Dairy Science 90, 4411–4419. doi:10.3168/jds.2007-0227

Mc Parland, S., Kearney, J.F., Rath, M., Berry, D.P., 2007b. Inbreeding trends and pedigree analysis of Irish dairy and beef cattle populations. Journal of Animal Science 85, 322–331. doi:10.2527/jas.2006-367

McGahern, A.M., Edwards, C.J., Bower, M.A., Heffernan, A., Park, S.D.E., Brophy, P.O., Bradley, D.G., MacHugh, D.E., Hill, E.W., 2006. Mitochondrial DNA sequence diversity in extant Irish horse populations and in ancient horses. Animal Genetics 37, 498–502. doi:10.1111/j.1365-2052.2006.01506.x

McHugo, G., Browett, S., Randhawa, I., Howard, D., Mullen, M., Richardson, I., Park, S., Magee, D., Scraggs, E., Dover, M., Correia, C., Hanrahan, J., MacHugh, D., 2019. A Population Genomics Analysis of the Native Irish Galway Sheep Breed. Frontiers in Genetics 10, 927. doi:10.3389/fgene.2019.00927

Meuwissen, T.H.E., Luo, Z., 1992. Computing inbreeding coefficients in large populations. Genetics, selection, evolution 24, 305–313. doi:10.1186/1297-9686-24-4-305

Nei, M., 1987. Molecular evolutionary genetics, 2. print. ed. Columbia Univ. Pr, New York, NY. Oldenbroek, J.K., 2017. Management of Animal Genetic Diversity. Wageningen Academic Publishers, Wageningen, the Netherlands.

Orozco-terWengel, P., Barbato, M., Nicolazzi, E., Biscarini, F., Milanesi, M., Davies, W., Williams, D., Stella, A., Ajmone-Marsan, P., Bruford, M.W., 2015. Revisiting demographic processes in cattle with genome-wide population genetic analysis. Frontiers in Genetics 6, 191. doi:10.3389/fgene.2015.00191

Park, K.-E., Kaucher, A.V., Powell, A., Waqas, M.S., Sandmaier, S.E.S., Oatley, M.J., Park, C.-H., Tibary, A., Donovan, D.M., Blomberg, L.A., Lillico, S.G., Whitelaw, C.B.A., Mileham, A., Telugu, B.P., Oatley, J.M., 2017. Generation of germline ablated male pigs by CRISPR/Cas9 editing of the NANOS2 gene. Scientific Reports 7, 40176. doi:10.1038/srep40176

Peach, J., 2019. The great British meadow. Royal Botanic Garden Kew. Retrieved on 8 September 2025 from https://www.kew.org/read-and-watch/great-british-meadow

Pérez-Pereira, N., Wang, J., Quesada, H., Caballero, A., 2022. Prediction of the minimum effective size of a population viable in the long term. Biodiversity and Conservation 31, 2763–2780. doi:10.1007/s10531-022-02456-z

Peter, C., Bruford, M., Perez, T., Dalamitra, S., Hewitt, G., Erhardt, G., 2007. Genetic diversity and subdivision of 57 European and Middle-Eastern sheep breeds. Animal Genetics 38, 37–44. doi:10.1111/j.1365-2052.2007.01561.x

Petersen, J.L., Mickelson, J.R., Rendahl, A.K., Valberg, S.J., Andersson, L.S., Axelsson, J., Bailey, E., Bannasch, D., Binns, M.M., Borges, A.S., Brama, P., da Câmara Machado, A., Capomaccio, S., Cappelli, K., Cothran, E.G., Distl, O., Fox-Clipsham, L., Graves, K.T., Guérin, G., Haase, B., Hasegawa, T., Hemmann, K., Hill, E.W., Leeb, T., Lindgren, G., Lohi, H., Lopes, M.S., McGivney, B.A., Mikko, S., Orr, N., Penedo, M.C.T., Piercy, R.J., Raekallio, M., Rieder, S., Røed, K.H., Swinburne, J., Tozaki, T., Vaudin, M., Wade, C.M., McCue, M.E., 2013. Genome-Wide Analysis Reveals Selection for Important Traits in Domestic Horse Breeds. PLoS Genetics 9, e54997. doi:10.1371/journal.pgen.1003211

Pocrnic, I., Lindgren, F., Tolhurst, D., Herring, W.O., Gorjanc, G., 2022. Optimisation of the core subset for the APY approximation of genomic relationships. Genetics Selection Evolution 54, 76. doi:10.1186/s12711-022-00767-x

Porter, V. (Ed.), 2002. Mason’s World Dictionary of Livestock Breeds, Types and Varieties. CABI Publishing, Wallingford, Oxon, UK.

Protected geographical food and drink names: UK GI schemes, 2024. UK Government. Retrieved on 8 September 2025 from https://www.gov.uk/guidance/protected-geographical-food-and-drink-names-uk-gi-schemes

Prystupa, J.M., Hind, P., Cothran, E.G., Plante, Y., 2012. Maternal Lineages in Native Canadian Equine Populations and Their Relationship to the Nordic and Mountain and Moorland Pony Breeds. Journal of Heredity 103, 380–390. doi:10.1093/jhered/ess003

Purcell, S., Neale, B., Todd-Brown, K., Thomas, L., Ferreira, M.A.R., Bender, D., Maller, J., Sklar, P., de Bakker, P.I.W., Daly, M.J., Sham, P.C., 2007. PLINK: A Tool Set for Whole-Genome Association and Population-Based Linkage Analyses. The American Journal of Human Genetics 81, 559–575. doi:10.1086/519795

Purfield, D.C., McParland, S., Wall, E., Berry, D.P., 2017. The distribution of runs of homozygosity and selection signatures in six commercial meat sheep breeds. PLOS ONE 12, e0176780. doi:10.1371/journal.pone.0176780

Rafter, P., McHugh, N., Pabiou, T., Berry, D.P., 2022. Inbreeding trends and genetic diversity in purebred sheep populations. animal 16, 100604. doi:10.1016/j.animal.2022.100604

Rauw, W.M., Kanis, E., Noordhuizen-Stassen, E.N., Grommers, F.J., 1998. Undesirable side effects of selection for high production efficiency in farm animals: a review. Livestock Production Science 56, 15–33. doi:10.1016/S0301-6226(98)00147-X

RBST, n.d. North Ronaldsay. Rare Breeds Survival Trust. Retrieved on 8 September 2025a from https://www.rbst.org.uk/north-ronaldsay

RBST, n.d. Shetland. Rare Breeds Survival Trust. Retrieved on 8 September 2025b from https://www.rbst.org.uk/shetland

RBST, n.d. Gloucestershire Old Spots. Rare Breeds Survival Trust. Retrieved on 8 September 2025c from https://www.rbst.org.uk/gloucestershire-old-spots

RBST, n.d. Shetland. Rare Breeds Survival Trust. Retrieved on 8 September 2025d from https://www.rbst.org.uk/shetland-geese

RBST, n.d. Scots Grey. Rare Breeds Survival Trust. Retrieved on 8 September 2025e from https://www.rbst.org.uk/scots-grey

RBST, n.d. Introduction. Rare Breeds Survival Trust. Retrieved on 8 September 2025f from https://www.rbst.org.uk/our-mission

RBST watchlist 2024, 2024. Rare Breeds Survival Trust (RBST). Retrieved on 8 September 2025 from https://www.rbst.org.uk/news/rbst-watchlist-2024

Rodríguez-Ramilo, S.T., Fernández, J., Toro, M.A., Hernández, D., Villanueva, B., 2015. Genome-Wide Estimates of Coancestry, Inbreeding and Effective Population Size in the Spanish Holstein Population. PLOS ONE 10, e0124157. doi:10.1371/journal.pone.0124157

Rohatgi, A., 2024. WebPlotDigitizer, v4. Retrieved on 8 September 2025 from https://automeris.io

Roughsedge, T., Brotherstone, S., Visscher, P.M., 1999. Quantifying genetic contributions to a dairy cattle population using pedigree analysis. Livestock Production Science 60, 359–369. doi:10.1016/S0301-6226(99)00106-2

Schwabe, A.E., Hall, S.J.G., 1989. Dystocia in nine British breeds of cattle and its relationship to dimensions of the dam and calf. Veterinary Record 125, 636– 639. doi:10.1136/vr.125.26-27.636

Stella, A., Nicolazzi, E.L., Van Tassell, C.P., Rothschild, M.F., Colli, L., Rosen, B.D., Sonstegard, T.S., Crepaldi, P., Tosser-Klopp, G., Joost, S., 2018. AdaptMap: exploring goat diversity and adaptation. Genetics Selection Evolution 50, 61, s12711–018-0427–5. doi:10.1186/s12711-018-0427-5

Tajima, F., 1983. EVOLUTIONARY RELATIONSHIP OF DNA SEQUENCES IN FINITE POPULATIONS. Genetics 105, 437–460. doi:10.1093/genetics/105.2.437

Taylor, L., Carlson, D.F., Nandi, S., Sherman, A., Fahrenkrug, S.C., McGrew, M.J., 2017. Efficient TALEN-mediated gene targeting of chicken primordial germ cells. Development 144, 928–934. doi:10.1242/dev.145367

Tenesa, A., Navarro, P., Hayes, B., Duffy, D., Clarke, G., Goddard, M., Visscher, P., 2007. Recent human effective population size estimated from linkage disequilibrium. GENOME RESEARCH 17, 520–526. doi:10.1101/gr.6023607

The Bovine HapMap Consortium, Gibbs, R.A., Taylor, J.F., Van Tassell, C.P., Barendse, W., Eversole, K.A., Gill, C.A., Green, R.D., Hamernik, D.L., Kappes, S.M., Lien, S., Matukumalli, L.K., McEwan, J.C., Nazareth, L.V., Schnabel, R.D., Weinstock, G.M., Wheeler, D.A., Ajmone-Marsan, P., Boettcher, P.J., Caetano, A.R., Garcia, J.F., Hanotte, O., Mariani, P., Skow, L.C., Sonstegard, T.S., Williams, J.L., Diallo, B., Hailemariam, L., Martinez, M.L., Morris, C.A., Silva, L.O.C., Spelman, R.J., Mulatu, W., Zhao, K., Abbey, C.A., Agaba, M., Araujo, F.R., Bunch, R.J., Burton, J., Gorni, C., Olivier, H., Harrison, B.E., Luff, B., Machado, M.A., Mwakaya, J., Plastow, G., Sim, W., Smith, T., Thomas, M.B., Valentini, A., Williams, P., Womack, J., Woolliams, J.A., Liu, Y., Qin, X., Worley, K.C., Gao, C., Jiang, H., Moore, S.S., Ren, Y., Song, X.-Z., Bustamante, C.D., Hernandez, R.D., Muzny, D.M., Patil, S., San Lucas, A., Fu, Q., Kent, M.P., Vega, R., Matukumalli, A., McWilliam, S., Sclep, G., Bryc, K., Choi, J., Gao, H., Grefenstette, J.J., Murdoch, B., Stella, A., Villa-Angulo, R., Wright, M., Aerts, J., Jann, O., Negrini, R., Goddard, M.E., Hayes, B.J., Bradley, D.G., Barbosa Da Silva, M., Lau, L.P.L., Liu, G.E., Lynn, D.J., Panzitta, F., Dodds, K.G., 2009. Genome-Wide Survey of SNP Variation Uncovers the Genetic Structure of Cattle Breeds. Science 324, 528–532. doi:10.1126/science.1167936

The Gloucestershire Old Spots Pig Breeders’ Club, 2006. Application for registration of a TSG council regulation (EC) No 509/2006 ‘Traditionally farmed Gloucestershire Old Spots Pork’ EC No UK/007-0024 TSG. Retrieved on 8 September 2025 from https://assets.publishing.service.gov.uk/media/5fd36838d3bf7f306291b5a3/pfn-traditionally-farmed-gloucs-old-spots-pork.pdf

The Great Trossachs Forest, 2025. Conservation grazing. The Great Trossachs Forest. Retrieved on 8 September 2025 from https://www.thegreattrossachsforest.co.uk/conservation-grazing

UN environment programme, 2022. Kumming-Montreal Global biodiversity framework (No. CBD/COP/15/L.25). Conference of the parties to the convention on biological diversity, Convention on Biological Diversity, Montreal, Canada. Retrieved on 8 September 2025 from https://www.cbd.int/doc/c/e6d3/cd1d/daf663719a03902a9b116c34/cop-15-l-25-en.pdf

United Nations General Assembly, 2015. Resolution adopted by the General Assembly on 25 September 2015: 70/1. Transforming our world: the 2030 Agenda for Sustainable Development. Retrieved on 8 September 2025 from https://docs.un.org/en/A/RES/70/1

Van Diepen, P., McLean, B., Frost, D., 2007. Livestock breeds and organic farming systems. ADAS Wales and Organic Centre Wales. Retrieved on 8 September 2025 from https://orgprints.org/id/eprint/10822/1/breeds07.pdf

Wall, E., Brotherstone, S., Kearney, J.F., Woolliams, J.A., Coffey, M.P., 2005. Impact of Nonadditive Genetic Effects in the Estimation of Breeding Values for Fertility and Correlated Traits. Journal of Dairy Science 88, 376–385. doi:10.3168/jds.S0022-0302(05)72697-7

Waples, R.S., 2025. The Idiot’s Guide to Effective Population Size. Molecular Ecology e17670. doi:10.1111/mec.17670

Webber, B.L., Raghu, S., Edwards, O.R., 2015. Is CRISPR-based gene drive a biocontrol silver bullet or global conservation threat? Proceedings of the National Academy of Sciences 112, 10565–10567. doi:10.1073/pnas.1514258112

Weigel, K.A., 2001. Controlling Inbreeding in Modern Breeding Programs. Journal of Dairy Science 84, E177–E184. doi:10.3168/jds.S0022-0302(01)70213-5

Weir, B.S., Cockerham, C.C., 1984. Estimating F-Statistics for the Analysis of Population Structure. Evolution 38, 1358–1370.

Weir, B.S., Hill, W.G., 1980. EFFECT OF MATING STRUCTURE ON VARIATION IN LINKAGE DISEQUILIBRIUM. Genetics 95, 477–488. doi:10.1093/genetics/95.2.477

Whittle, A.W.R., Healy, F., Bayliss, A., Allen, M.J., 2011. Gathering timeL: dating the early Neolithic enclosures of southern Britain and Ireland. Oxbow Books, Oxford.

Wiener, P., Burton, D., Williams, J.L., 2004. Breed relationships and definition in British cattle: a genetic analysis. Heredity 93, 597–602. doi:10.1038/sj.hdy.6800566

Wilkinson, S., Wiener, P., Teverson, D., Haley, C.S., Hocking, P.M., 2012. Characterization of the genetic diversity, structure and admixture of British chicken breeds. Animal Genetics 43, 552–563. doi:10.1111/j.1365-2052.2011.02296.x

Williams, J.L., Hall, S.J.G., Del Corvo, M., Ballingall, K.T., Colli, L., Ajmone Marsan, P., Biscarini, F., 2015. Inbreeding and purging at the genomic Level: the Chillingham cattle reveal extensive, non_-_random SNP heterozygosity. Animal Genetics 47, 19–27. doi:10.1111/age.12376

Winton, C.L., Hegarty, M.J., McMahon, R., Slavov, G.T., McEwan, N.R., Davies_-_Morel, M.C.G., Morgan, C.M., Powell, W., Nash, D.M., 2013. Genetic diversity and phylogenetic analysis of native mountain ponies of Britain and Ireland reveals a novel rare population. Ecology and Evolution 3, 934–947. doi:10.1002/ece3.507

Winton, C.L., McMahon, R., Hegarty, M.J., McEwan, N.R., Davies_-_Morel, M.C.G., Morgan, C., Nash, D.M., 2020. Genetic diversity within and between British and Irish breeds: The maternal and paternal history of native ponies. Ecology and Evolution 10, 1352–1367. doi:10.1002/ece3.5989

Wright, S., 1950. THE GENETICAL STRUCTURE OF POPULATIONS. Annals of Eugenics 15, 323–354. doi:10.1111/j.1469-1809.1949.tb02451.x

Wright, S., 1931. Evolution in Mendelian populations. Genetics 16, 97–159. doi:10.1093/genetics/16.2.97

Wright, S., McPhee, H.C., 1925. An approximate method of calculating coefficients of inbreeding and relationship from livestock pedigrees. Journal of Agricultural Research 31, 377–383.

Wykes, D.L., 2004. Robert Bakewell (1725–1795) of Dishley: farmer and livestock improver. The Agricultural History Review 52, 38–55.

Yu, X., Woolliams, J.A., Meuwissen, T.H., 2014. Prioritizing animals for dense genotyping in order to impute missing genotypes of sparsely genotyped animals. Genetics Selection Evolution 46, 46. doi:10.1186/1297-9686-46-46

Zeder, M. a, 2008. Domestication and early agriculture in the Mediterranean Basin: Origins, diffusion, and impact. Proceedings of the National Academy of Sciences of the United States of America 105, 11597–11604. doi:10.1073/pnas.0801317105

